# Aphid feeding induces the relaxation of epigenetic control and the associated regulation of the defense response in *Arabidopsis*

**DOI:** 10.1101/2020.01.24.916783

**Authors:** Maria Luz Annacondia, Dimitrije Markovic, Juan Luis Reig-Valiente, Vassilis Scaltsoyiannes, Corné M.J. Pieterse, Velemir Ninkovic, R. Keith Slotkin, German Martinez

## Abstract

Environmentally induced changes in the epigenome help individuals to quickly adapt to fluctuations in the conditions of their habitats. Here we explored those changes in *Arabidopsis thaliana* plants subjected to multiple biotic and abiotic stresses, and identified transposable element (TE) activation in plants infested with the green peach aphid, *Myzus persicae*. We performed a genome-wide analysis of DNA methylation, mRNA expression, mRNA degradation and small RNA accumulation. Our results demonstrate that aphid feeding induces loss of methylation of hundreds of loci, mainly TEs. This loss of methylation has the potential to regulate gene expression and we found evidence that it is involved in the control of key plant immunity genes. Accordingly, we find that mutant plants deficient in epigenetic silencing show increased resistance to *M.persicae* infestation. Collectively, our results show that changes in DNA methylation play a significant role in the regulation of the plant transcriptional response and induction of defense response against aphid feeding.

## INTRODUCTION

While adaptation to long-term environmental changes involves genetic variation, fluctuating stresses are normally coped through the modulation of the transcription machinery (Lamke & Baurle, 2017). Several mechanisms govern the transcriptional response during stress including transcription factors (TFs) and epigenetic regulation (Gutzat & Mittelsten Scheid, 2012). In eukaryotic organisms epigenetic modifications of chromatin and DNA are the core of genome stability regulation through the control of transposable element (TE) expression and transposition (Law & Jacobsen, 2010). Epigenetic modifications consist of covalent and reversible marks that are deposited on both the DNA and the histones. DNA methylation constitutes a vital and widespread mark in plant genomes, where it can happened in three different sequence combinations: the symmetric contexts CG and CHG, and the asymmetric CHH (where H can be A, C or T) (Law & Jacobsen, 2010). This repressive mark is established by the action of small RNAs (sRNAs) through a pathway named RNA-directed DNA methylation (RdDM) and can be actively removed from any context by the action of DNA glycosylases (Matzke & Mosher, 2014, Zhang, Lang et al., 2018). The modifications that occur in the tails of histones can be active or repressive marks. For example, H3K4 mono-, di- and tri-methylation (H3K4me1, H3K4me2 and H3K4me3) are associated with highly transcribed genes (Zhang, Bernatavichute et al., 2009), H3K27 tri-methylation (H3K27me3) and is mainly found in silenced genes (Zhang, Clarenz et al., 2007) and H3K9 di-methylation (H3K9me2) is rarely seen in genes while is predominantly present in TEs, where it correlates with the presence of DNA methylation, leading to transcriptional silencing and the formation of heterochromatin (Zhou, Wang et al., 2010).

TEs are a source of new mutations and genetic/genomic variation and, interestingly, of new regulatory regions for genes (Kidwell & Lisch, 1997, Lisch, 2009). Several agricultural traits like orange, maize and apple color or pepper pungency are regulated by TEs that inserted in new locations and created new expression patterns for the gene/s in the vicinity of the insertion (Butelli, Licciardello et al., 2012, Dooner, Robbins et al., 1991, Tanaka, Asano et al., 2019, Zhang, Hu et al., 2019). These TE domestication events are especially important for plant interaction with their environment (Annacondia, Mageroy et al., 2018). Different abiotic and biotic stresses (including drought, salinity, heat, cold, ultraviolet radiation, chemical agents and viral, viroid, bacterial and fungal infections) show examples of TE domestication events that influence gene expression and/or induce changes in the epigenetic regulation of repeats (Annacondia et al., 2018, Mozgova, Mikulski et al., 2019). Defense genes are interesting examples of the interaction between epigenetic regulation and gene regulation and evolution, since most nucleotide binding site and leucine-rich repeat domain protein (NBS-LRR) genes accumulate in heterochromatic clusters populated by TEs (Meyers, Kozik et al., 2003). As an example of the role of epigenetic regulation in their transcriptional control, several defense genes such as *RECOGNITION OF PERONOSPORA PARASITICA 7* (*RPP7*), *RPP4* and *RESISTANCE METHYLATED GENE 1* (*RMG1*) are transcriptionally regulated by domesticated TEs (Tsuchiya & Eulgem, 2013, Yu, Lepere et al., 2013, Zervudacki, Yu et al., 2018). Additionally, mutants of different DNA methylation, RdDM and small RNA pathways regulate immunity to bacterial and fungal infection (Agorio & Vera, 2007, Dowen, Pelizzola et al., 2012, Lopez, Ramirez et al., 2011, Yu et al., 2013). Intriguingly, some biotic stresses can induce tolerance towards the pathogen in the subsequent generation (Boyko, Blevins et al., 2010, Boyko, Kathiria et al., 2007, De Vos & Jander, 2009, Kathiria, Sidler et al., 2010, Luna, Bruce et al., 2012, Slaughter, Daniel et al., 2012), a phenomenon that could be explained by changes in the methylation status of the DNA or chromatin rather than by spontaneous mutagenesis and reversion (Annacondia & Martinez, 2019, Boyko & Kovalchuk, 2011, Luna & Ton, 2012).

The relationship between pathogens and host plants involves an interaction between both genomes and leads to events of coevolution. An example of this interaction takes place between plants and insects. Both groups interact in different ways and have influenced each other during evolution (e.g. the appearance in land plants of entomophily (Darwin, 1899) or carnivory (Renner & Specht, 2013) or the artificial selection of insects that evolve resistance to plants with defense genes (Bown, Wilkinson et al., 1997). Plant-insect interactions are classified as mutualistic, antagonistic or commensalistic. Although they are basic for the ecological equilibrium, some of them can be a threat for the agricultural ecosystems and, by hence, to food production. Herbivory insects represent approximately 50% of the total insect species (Schoonhoven, van Loon et al., 2005) and are considered a threat to plant productivity. They are among the stresses that induce trans-generational acquired resistance, pointing to a potential role of epigenetic regulation of plant defense (De Vos & Jander, 2009, Rasmann, De Vos et al., 2012). Nevertheless, how this epigenetic response is established during insect infestation is poorly characterized.

Here, we report that epigenetic control is an important part of the *Arabidopsis thaliana* defense response against the infestation by the green peach aphid *Myzus persicae*. Our analysis of DNA methylation, mRNA, small RNAs and mRNA cleavage changes induced in plants exposed to aphid feeding shows that the response of the plant is characterized by a transcriptional reprogramming and methylation changes in TEs. These TEs are normally associated with repressive/heterochromatic marks and dependent on the RdDM pathway for their silencing. Along with this, we find that upon infestation certain differentially methylated regions (DMRs) are associated with infestation-responsive genes and TF binding sites. Finally, we find that mutant plants deficient in epigenetic silencing show increased resistance to *M.persicae* infestation. Together, our data uncovers a novel role of plant epigenetic control in the induction of the transcriptional response to aphid feeding.

## RESULTS

### Meta analysis of TE activation identifies *Myzus persicae* as a potential inducer of epigenetic changes

To identify stresses that alter the epigenetic regulation in *Arabidopsis thaliana*, we performed a meta-analysis of TE expression from ATH1 microarray datasets, which have been widely used by the community. The ATH1 microarray contains 1155 TE probes used to track changes in transcript abundance influenced by epigenetic reprogramming (Slotkin, Vaughn et al., 2009). We investigated TE expression under different stresses including abiotic (heavy metal presence, exposure to heat, cold, spaceflight or UV light among others) and biotic (viral, oomycete, bacterial and insect infection/infestation) (Figure 1A, Supplementary Table 1 and data not shown). We found that, in general, these stresses can induce a modest reactivation of TEs, although this response is dependent on the specific stress (Figure 1A and B). Interestingly, biotic stress seems to activate TE expression more consistently than the abiotic stresses analyzed here (Figure 1A-B). This analysis identified that among the stresses inducing TE reactivation, *M.persicae* infestation after 72 hrs induces the highest TE transcription. *M.persicae* is a major agricultural pest to a large variety of plants that include stone fruits, potato and horticultural crops (Louis & Shah, 2013). A high number of TEs (533 TEs, 46.1% of all the TEs represented in the ATH1 microarray, Figure 1B) show evidence of transcriptional activation when plants were under attack from *M.persicae* compared to control plants. This reactivation included more than 40% of all the DNA transposons and retrotransposons represented in the ATH1 microarray and is enriched in *Gypsy* and *Copia* retrotransposons and TIR DNA transposons (Figure 1C). Analysis of the reactivation indicated that TE activation takes place at 48 hrs and increases by 72 hrs pi (Supplementary Figure 1, average fold change value for retrotransposons at 72 hrs pi 3.64x and 4.4x for DNA transposons). Other cases of largescale TE activation are seen when DNA methylation, histone modification and/or heterochromatin formation are lost (Lippman, Gendrel et al., 2004, Lippman, May et al., 2003, Panda, Ji et al., 2016, Zilberman, Gehring et al., 2007). Together, these results indicate that *M.persicae* infestation results in TE activation, potentially due to a large-scale change in the epigenome.

**Figure 1.**
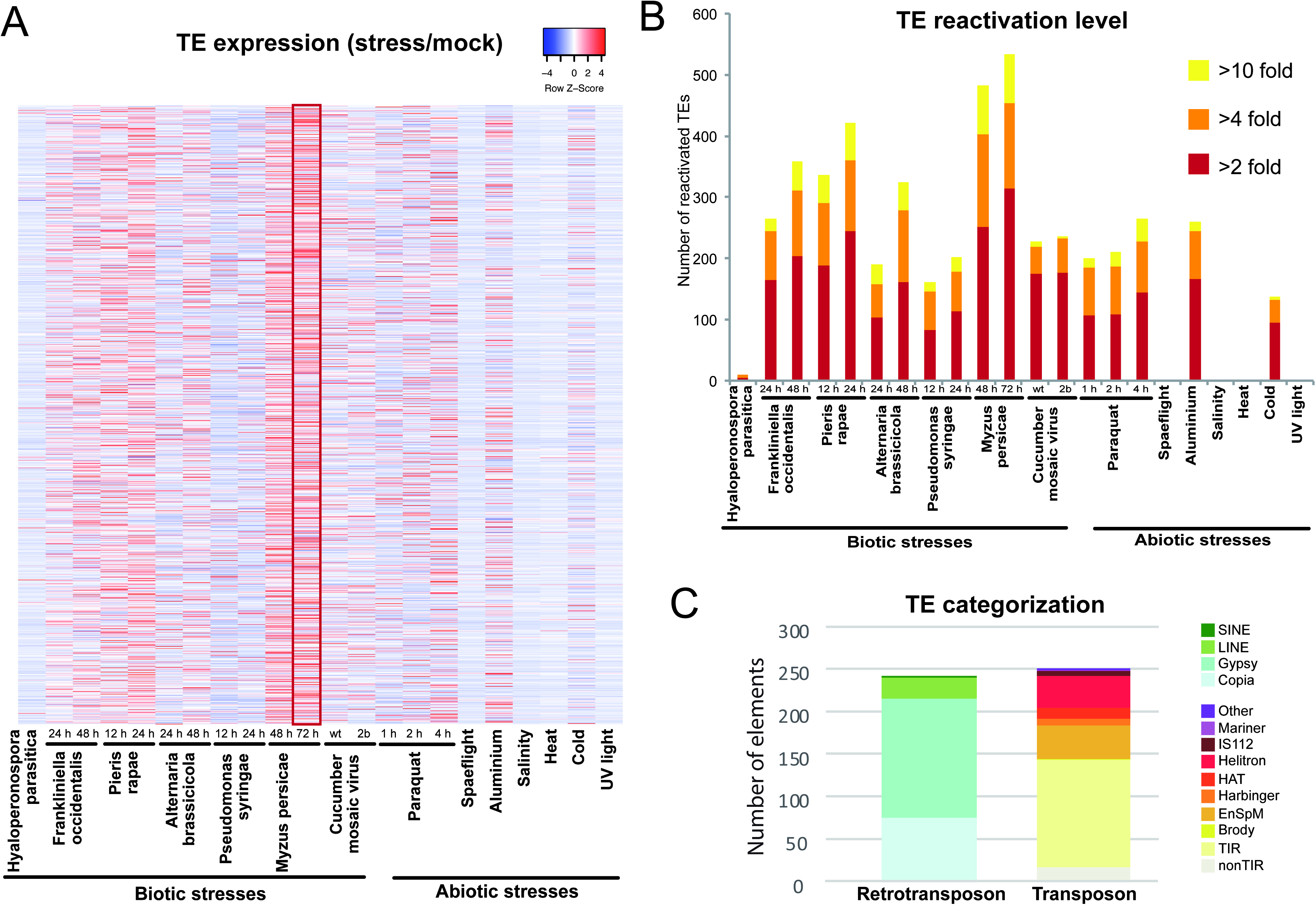
*M.persicae* infestation induces TE reactivation. A. Meta-analysis of TE expression in the ATH1 microarray in several stresses. Heat map of the expression values of the indicated treatment relative to their respective control. In experiments with several bioreplicates the mean values between bioreplicates was used. B. Number of TEs reactivated in the analyzed stresses grouped by fold categories. C. Percentage of reactivated TEs belonging to different categories in the ATH1 microarray.

### Transcriptional response to aphid feeding in *Arabidopsis* is characterized by transcription factor activity

The extent of TE reactivation observed in our meta analysis could be biased by the presence of TE probes on the ATH1 microarray. To monitor the transcriptional changes under aphid infestation, we repeated the experiment described in (De Vos, Van Oosten et al., 2005) and analyzed in Figure 1 at 72 hrs post infestation (p.i.) and prepared and sequenced high-throughput mRNA libraries (Supplementary Table 2). First, we focused on understanding the genic transcriptional changes taking place in Arabidopsis infested with *M.persicae*. This analysis revealed that 267 genes are significantly differentially expressed, with almost all of these being upregulated (265 genes, Figure 2A and Supplementary Table 2). As expected, the analysis of the GO categories for significantly upregulated genes indicates that these genes are associated with the response to stress or environmental stimuli (Figure 2B and C and Supplementary Figure 2A). Interestingly differentially expressed genes contain a significant overrepresentation of mobile mRNAs (24.34% of differentially expressed genes, two tailed p<0.0001 calculated by a Chi-squared test with Yates correction) (Thieme, Rojas-Triana et al., 2015) (Supplementary Figure 2B).

**Figure 2.**
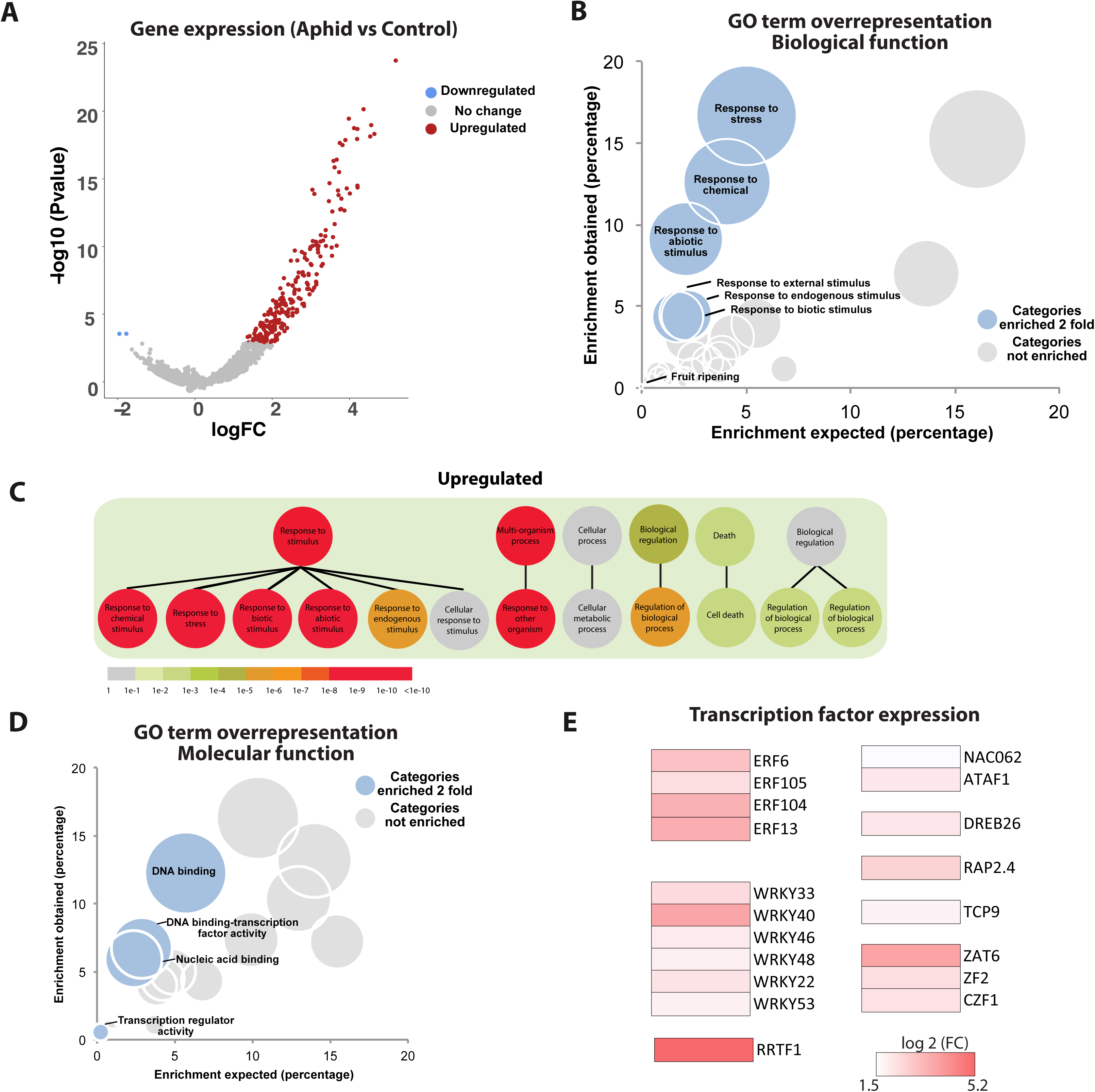
Aphid feeding-induced changes in gene expression. A. Volcano plot depicting gene expression in the comparison aphid vs control sample. Dots colored in red indicated genes with significant upregulation. B. Bubble graph depicting the GO term overrepresentation test for upregulated genes grouped by biological function. Bubbles in blue show GO categories enriched two fold or more. C. Biomap of upregulated genes. D. Bubble graph depicting the GO term overrepresentation test for upregulated genes grouped by molecular function. Bubbles in blue show GO categories enriched two fold or more. E. Examples of different transcription factors showing upregulation during aphid infestation.

We further analyzed the molecular functions of these stress-responsive genes by checking the GO term enrichment according to molecular function (Figure 2D). This revealed an overrepresentation of DNA binding/transcription factor categories, indicating that these transcriptional regulators are an important part of the response to aphid feeding (Figure 2D). Interestingly, several well-studied TFs show a strong upregulation (higher than 1.5 log2 fold change) including several members of the WRKY and ERF families (Figure 2E), which have been previously associated with the response against aphid feeding (Gao, Kamphuis et al., 2010). In summary, the transcriptional response against aphids shows an overrepresentation of TF activity.

### Complex transcriptional and posttranscriptional regulation of TEs during aphid infestation

Our previous analysis of ATH1 public datasets indicated a potential reactivation of TEs during aphid infestation. However, the TE probes on the ATH1 array do not represent the genomic distribution of TEs, and favor *Helitron* elements that resemble genes. Accordingly, we explored TE transcriptional and posttranscriptional regulation by performing RNA, PARE and sRNA sequencing, which target (respectively) mRNAs, mRNAs targeted for degradation and sRNAs derived from Pol II and Pol IV activity (Figure 3). For the production of PARE libraries we used the same tissue as in our mRNA analysis, with plants infested with *M.persicae* for 72hrs. Analysis of RNA sequencing indicated that, surprisingly, TE changes at the steady-state mRNA level, if any, are minimal (Figure 3A). This difference between the microarray and mRNA-seq data may be due to the ability to map sequencing reads to TEs and/or the degradation of TE mRNAs that will still be detected by 3’ microarray probes. On the other hand, PARE sequencing was able to identify changes of TE expression that experience uncharacterized posttranscriptional regulation (Figure 3B). We identified 73 TEs that increase their transcription during aphid infestation while 42 are downregulated (Figure 3B). This apparently contradictory output from RNA and PARE sequencing indicates that there is indeed transcriptional reactivation of TEs, but these transcripts are regulated at the posttranscriptional level since they are only detectable by PARE sequencing, which specifically captures mRNAs with a 5’ P that are degradation intermediates (Addo-Quaye, Eshoo et al., 2008, German, Pillay et al., 2008, Hou, Lee et al., 2016, Pelechano, Wei et al., 2015, Yu, Willmann et al., 2016).

**Figure 3.**
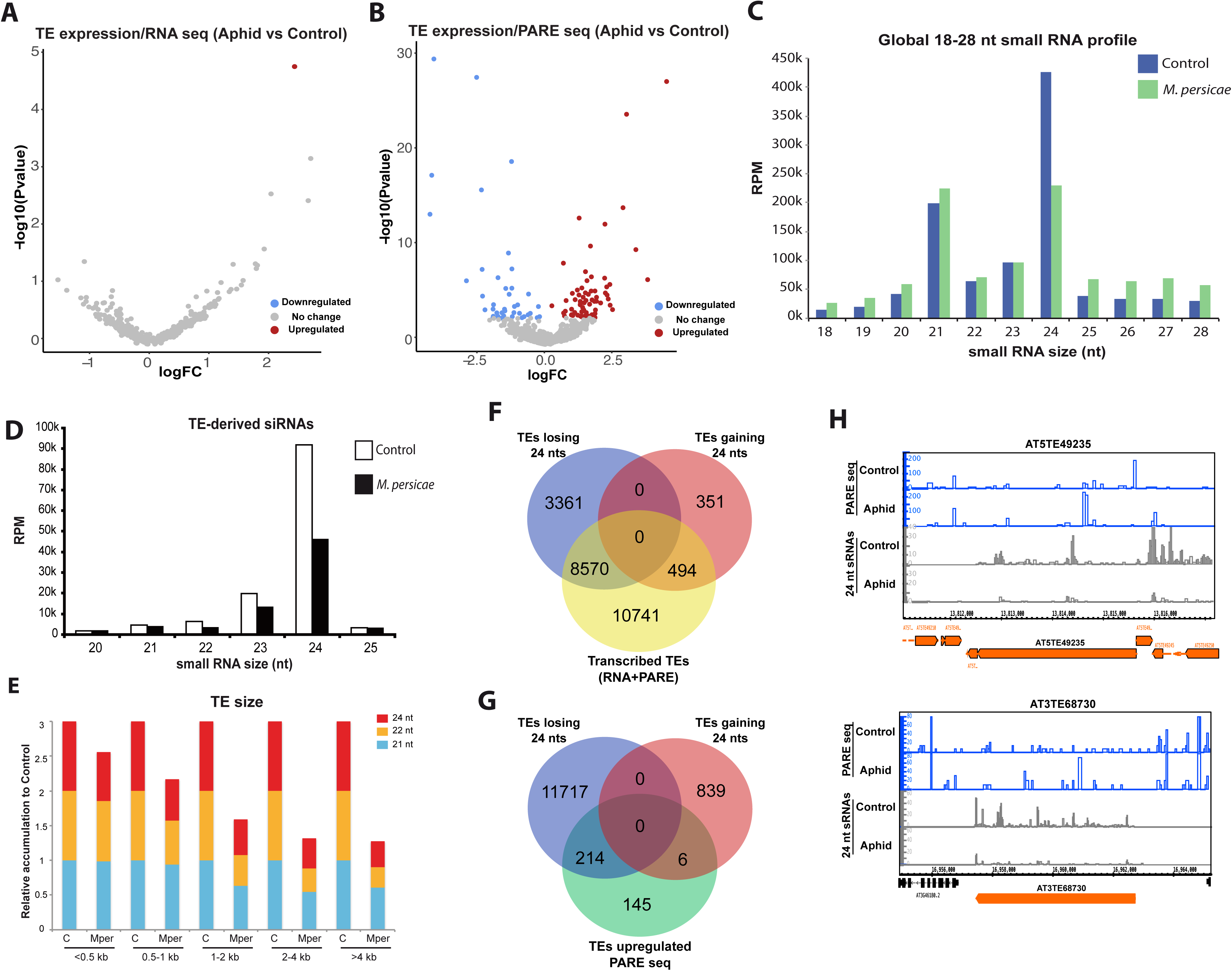
Changes induced at TE expression by aphid feeding. A. Volcano plot depicting TE mRNA-seq expression in the comparison aphid vs control RNA samples. Dots colored in red indicated genes with significant upregulation. B. Volcano plot depicting gene expression in the comparison aphid vs control PARE samples. Dots colored in red indicated genes with significant upregulation. C. Global sRNAs profiles of control and stressed samples. D. TE-derived sRNA profiles of control and stressed samples. E. Relative accumulation of 21,22 and 24 nt sRNAs in control (C) and aphid infested samples (Mper) for TEs of different sizes. Values shown are relative to control, where accumulation values for each sRNA category were set to 1. F. Venn diagram showing the overlap between the TE populations identified from each of the different RNA sequencing analyses. G. Venn diagram depicting the overlap of TEs upregulated two fold in the PARE sequencing data and TEs losing or gaining two fold 24 nt sRNAs. G. Screenshot of a genome browser showing the accumulation of PARE reads and 24 nt sRNAs in control and aphid samples for two of the TEs upregulated in the PARE libraries and showing a decrease of 24 nt sRNA accumulation.

Next, the analysis of our sRNA sequencing revealed more dramatic differences taking place almost exclusively at 24 nt TE-derived sRNAs (Figure 3C-D and Supplementary Figure 3B-H). This loss of 24 nt sRNAs is more pronounced on long transposons of the *Gypsy*, *Copia*, *MuDR* and *LINE* families (Figure 3E and Supplementary Figure 3B). Long retrotransposons are located in centromeric and pericentromeric regions, which are the genomic habitats of *Gypsy* and *Copia/LINE* elements, respectively (Underwood, Henderson et al., 2017). Altogether, this indicates that the loss of RdDM activity under aphid feeding takes place mainly at centromeric and pericentromeric regions.

Lastly, we analyzed the connection between the changes observed at the transcriptional (RNA and PARE sequencing) and sRNA level. First, 43.27% of TEs with evidence of transcription (TEs with reads in either RNA or PARE sequencing libraries) are associated with loss of 24 nt sRNAs (Figure 3F). Most of these TEs are detected by PARE sequencing (Supplementary Figure 3A), which might indicate that TE mRNA degradation could be associated with alternative pathways of 24 nt biogenesis. Nevertheless, TEs detected as upregulated by PARE sequencing are mostly associated with loss of 24 nt sRNAs (Figure 3G, example shown in Figure 3H). This fact, together with the lack of evidence for production of 24 nt sRNAs from polyadenylated transcripts during aphid infestation favors the hypothesis that loss of 24 nt sRNAs causes the transcriptional upregulation of TEs. In summary, our RNA, PARE and sRNA sequencing data indicates that during aphid infestation plants reduce the activity of the RdDM pathway, leading to the transcriptional reactivation of TEs that are, in turn, regulated at the posttranscriptional level.

### Differential methylation of the Arabidopsis genome upon aphid infestation

The transcriptional changes observed and the loss of TE-derived 24 nt sRNAs lead us to analyze the levels of DNA methylation. Genomic DNA was isolated, treated with sodium bisulfite and sequenced at 26.4 x average coverage (Supplementary Table 2). The data was plotted as a heat map on all five chromosomes comparing the control and aphid infested samples (Figure 4A). This data reveals a strong enrichment of DNA methylation in the pericentromeric heterochromatin, as expected from somatic tissues. A global analysis of the methylation level at genes and TEs for each methylation context revealed that, overall, no dramatic differences exist between the control and aphid infested samples in the overall profiles (Figure 4B). This is expected, since aphids cause very subtle wounding due to their feeding strategy.

**Figure 4.**
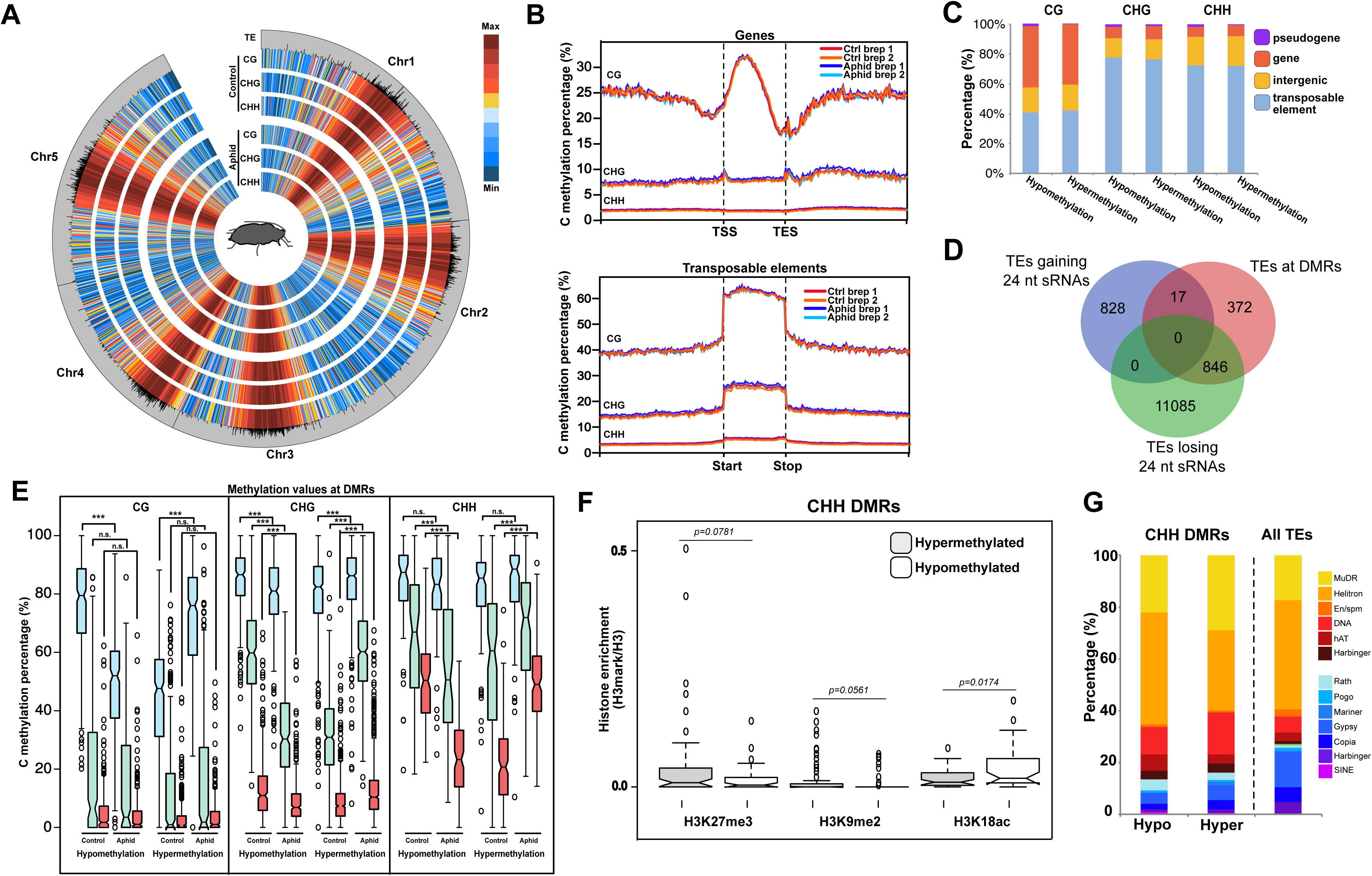
DNA methylation changes induced by aphid feeding. A. Genome-wide methylation levels for each of the C methylation contexts (CG, CHG and CHH) in control and aphid infested samples. B. DNA methylation coverage for genes and TEs for each C methylation context. C. Hypermethylation and hypomethylation DMRs identified for each C methylation context. D. DMR co-localization with different genomic entities. E. Cytosine methylation values at hypermethylation and hypomethylation DMRs for each methylation context. Asterisks indicate the different levels of significance between the comparisons (*<0.05, **<0.01, ***<0.001). p-value was calculated using an unpaired t-test. F. H3K27me3, H3K9me2 and H3K18ac enrichment relative to H3 for hypermethylated and hypomethylated DMRs. p-values were calculated using an unpaired t-test. G. Categorization of TEs co-localizing with CHH hypermethylation and hypomethylation DMRs in comparison to all the TEs in the TAIR10 *Arabidopsis* genome.

To identify regions in the genome harboring differential methylation upon aphid feeding we determined differentially methylated regions (DMRs) (Catoni, Tsang et al., 2018). This analysis revealed the presence of DMRs for all the DNA methylation contexts and associated both with hypo- and hypermethylation (Supplementary Figure 4A and Figure 4E). The CHG context has the greatest amount of DMRs (1123) followed by CG (691) and CHH (311). Furthermore, while CG DMRs are both present at genes and TEs, most of CHG and CHH DMRs are associated with TEs (Figure 4C). Interestingly, TEs located at DMRs are mostly the same TEs that lose 24 nt sRNAs (Figure 4D). DMRs in the CG context have low CHG and CHH methylation values and the changes experienced during aphid feeding in these contexts are not significant (Figure 4E), pointing to their association with gene body methylation (Figure 4C). CHG and CHH DMRs on the other hand are highly dynamic and experience significant changes in other methylation contexts (especially in the CHG and CHH contexts) in the regions that experience hypo and hypermethylation (Figure 4E). Due to the tight association between CHG and CHH methylation with H3K9me2 (Du, Johnson et al., 2015), this might indicate that a strong reorganization of heterochromatin takes place in these regions upon aphid feeding.

The relatively low number of DMRs and the lack of overall changes in the global profiles of DNA methylation may indicate that methylation changes only take place in specific regions. To test if DMRs might be associated with particular histone marks, we retrieved public datasets of different histone modifications coverage in *Arabidopsis* somatic tissues (Luo, Sidote et al., 2013) and checked the enrichment of those histone marks in our identified DMRs. Hypomethylated DMRs in the CHH context show enrichment in the permissive mark H3K18ac, while showing low amounts of the repressive marks H3K27me3 and H3K9me2 when compared to hypermethylated DMRs (Figure 4F and Supplementary Figure 4B-D). This indicates that removal of CHH methylation during aphid infestation only takes place at regions of the genome that have a high level of permissive histone marks and a low level of repressive histone marks. Furthermore, hypomethylated CHH DMRs show an enrichment in *Helitron* elements (two tailed p=0.0045 calculated by a Chi-squared test with Yates correction compared to presence of *Helitron* elements in the whole genome, Figure 4G), which are known to locate in the proximities of genes and influence their expression (Underwood et al., 2017). Therefore, upon aphid feeding, very localized methylation changes take place mainly associated with epigenetic labile TE regions.

### Stress-induced changes in methylation are associated with expression changes in defense-associated genes

Changes in TE methylation can influence the expression of neighboring genes (Wang, Weigel et al., 2013). To test if this occurs during aphid feeding, we obtained the list of neighbor genes within a 4kb window (2 kb upstream and downstream) for each DMR. This strategy identified 1010 genes associated with hypermethylated DMRs and 661 genes associated with hypomethylated DMRs (Supplementary Table 4). Since hypomethylation is expected to affect gene expression we focused our analysis on this category. Genes located in the proximities of hypomethylated DMRs are associated with oxygen binding, translation regulator activity, nuclease and motor activity, and fruit ripening and cell death when associated by biological function (>1.5 fold enrichment, Supplementary Figure 5A-B). When the GO categories are restrained to genes that show a significant change of expression (16 genes), we obtained an enrichment in genes with protein biding activity functions, and fruit ripening, cell death, pollination; and response to endogenous, chemical, external and biotic stimulus when grouped by biological function (>2 fold enrichment, Figure 5A). The partial lack of a higher number of genes with significant changes in expression associated with hypomethylated DMRs indicates that the presence of a hypomethylated DMR is not a condition to induce significant changes in gene expression *per se.* Probably, other regulatory elements are needed to reprogram the transcriptional response to aphid feeding.

**Figure 5.**
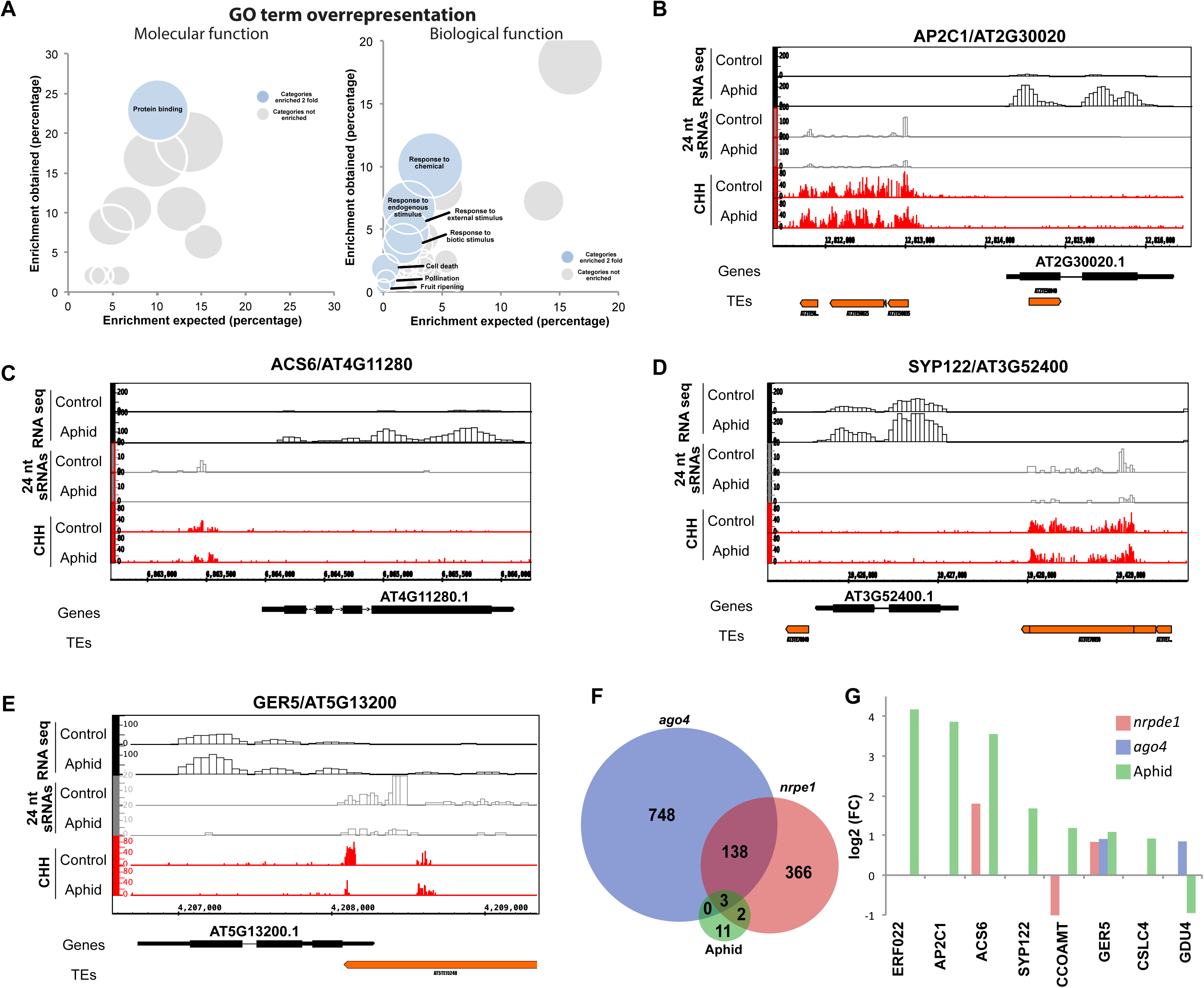
Transcriptional changes associated with DMRs. A Bubble graph depicting the GO term overrepresentation test for upregulated genes grouped by molecular (left panel) or biological function (right panel). Bubbles in blue show GO categories enriched two fold or more. B-E Examples of upregulated genes associated with CHH DMRs. F. Venn diagram depicting overlap between differentially and significant expressed genes in *pol v*, *ago4* and DMR associated genes in aphid infested samples. G. Expression of CHH associated differentially expressed genes in *pol v*, *ago4* and aphid infested samples. Only values from significant differences are shown (p-value<0.05).

We identified several significantly overexpressed genes located in the proximity of CHH hypomethylated DMRs that are related to plant defense (Figure 5B-E). These genes include *AP2C1*, a PP2C-type phosphatase that modulates innate immunity (Schweighofer, Kazanaviciute et al., 2007), *ACS6*, a 1-aminocyclopropane-1-carboxylic acid synthase a rate-limiting enzyme that catalyses the committing step of ethylene biosynthesis (Joo, Liu et al., 2008), *SYP122*, a Qa-SNARE proteins that drive vesicle fusion and are important for cell growth and expansion and pathogen defense (Waghmare, Lileikyte et al., 2018), *GER5* an stress-responsive glucosyltransferases, rab-like GTPase activators and myotubularin domain protein involved in ABA-mediated stress responses (Baron, Schroeder et al., 2014), the ethylene response factor *ERF022* and the caffeoyl-CoA 3-O-methyltransferase *CCOAMT* involved in the lignin biosynthesis pathway the accumulation of which induces resistance to aphid feeding (Wang, Sheng et al., 2017).

Next, we explored whether the expression of these genes was also altered in epigenetic mutants (not during aphid feeding). We used public data from Pol V and AGO4 mutants (Rowley, Rothi et al., 2017, Zhu, Rowley et al., 2013). Pol V and AGO4 are components of the RdDM pathway that produces sRNAs to target genomic regions and introduces DNA methylation (Matzke & Mosher, 2014). Pol V produces long non-coding transcripts that guide Pol IV-derived 24 nt sRNAs loaded into AGO4 to chromatin (Wierzbicki, Ream et al., 2009). Mutations in AGO4 or PolV impair RdDM-dependent methylation especially in the CHH context, and indeed 82% of loci regulated by Pol V or Pol IV are also regulated by AGO4/AGO6 (Duan, Zhang et al., 2015). Differentially expressed genes associated with DMRs are significantly enriched in genes regulated by the RdDM pathway components AGO4 and/or Pol V (31.25% overlap, two tailed p<0.0001 calculated by a Chi-squared test with Yates correction, Figure 5F-G). Interestingly, although some genes show a similar expression pattern between the RdDM mutants and the aphid-infested samples (e.g. *GER5*, *ACS6*, Figure 5G) others show opposing patterns of expression between the aphid infested samples and the RdDM mutants (notably *CCOAMT* and *GDU4*). This different expression pattern led us to question if the expression of these genes could be regulated by TFs that are not overexpressed in the RdDM mutants. Interestingly, the analysis of TF binding motifs present in the DMRs of differentially expressed genes associated with loss of CHH methylation showed several highly enriched including B3 binding domain-containing TFs like B3/ARF, AP2/B3 and B3 (20.65, 11.8 and 8.6 fold enrichment respectively, Supplementary Figure 5C and E). Several TFs of these families are differentially expressed in the aphid infested samples, while they do not show this pattern of expression in RdDM mutants (Supplementary Figure 5D). This indicates that differential expression of TFs likely leads to the observed differences in the expression pattern between aphid infested samples and RdDM mutants. Overall, our data indicates that DNA methylation changes are associated with gene expression changes, likely in combination with TF-induced expression.

### Epigenetic mutants show enhanced defense against aphids

Finally, we tested whether different *Arabidopsis* mutants defective in epigenetic regulation are more resistant to aphid infestation. For this, we analyzed aphid no-choice settling where 10 aphids were transferred to a random caged leaf (Figure 6A). We performed this test in different mutants including the histone remodeler *DDM1*, the triple mutant defective in maintenance of non-CG methylation *ddc* (*drm1 drm2 cmt3*), the main subunit of the principal factor of the RdDM pathway RNA Pol IV (*nrpd1*) and the H3K9me2 methyltransferase *KYP* (Figure 6B). Our analysis indicated that, from these components, mutations in *nrpd1* (the largest subunit of Pol IV) and *kyp* show a reduced number of aphids settled, and only *kyp* had a significant decrease (Figure 6B).

**Figure 6.**
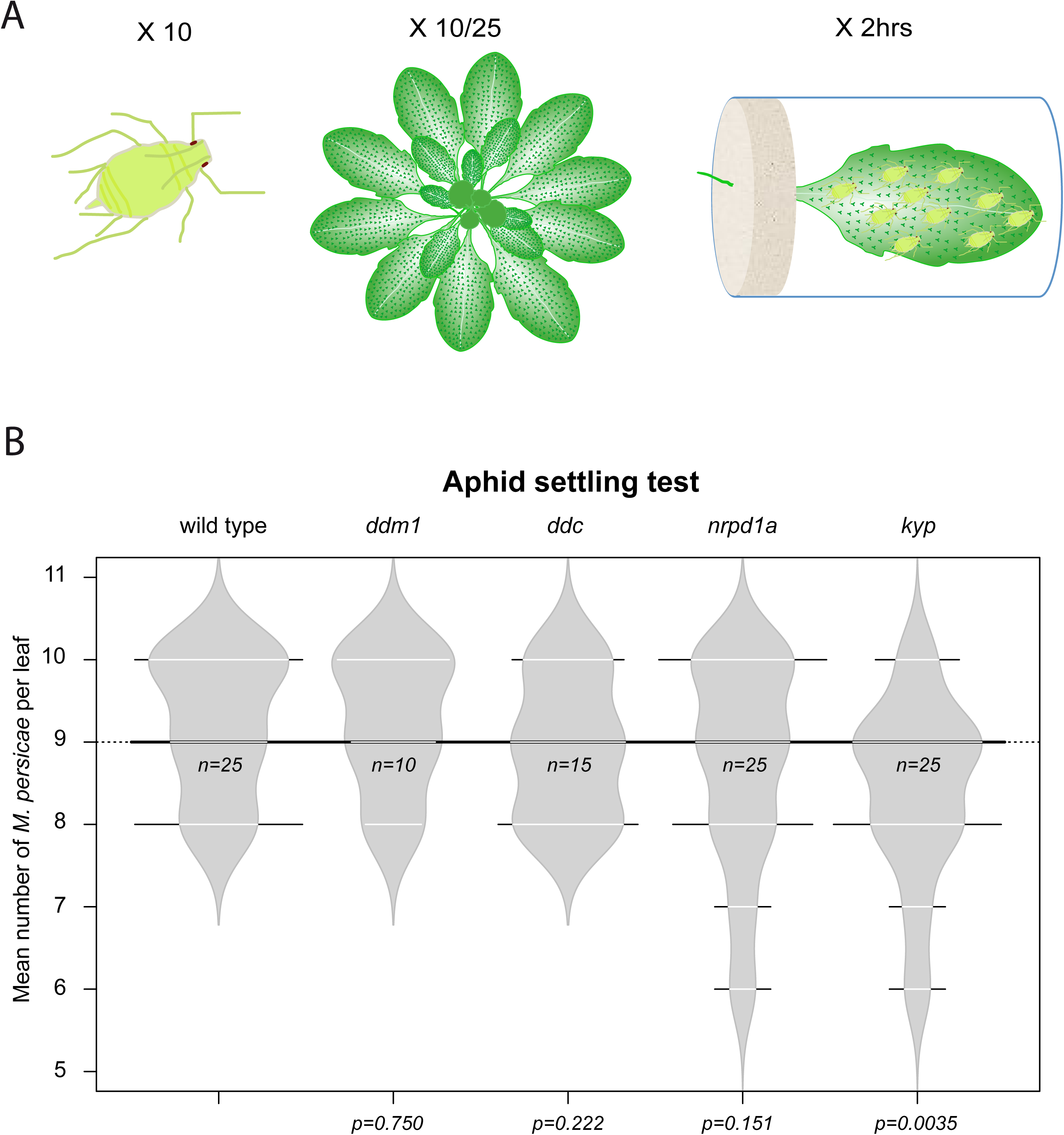
Epigenetic mutants are resistant to aphid settlement. A. Depiction of the aphid settlement experiment carried out in our analysis. In brief, 10 aphids were moved to a single caged leaf (attached to the plant) from 10/25 individual Arabidopsis plants. B. Aphid settlement test in different epigenetic mutants. p-values shown were calculated using an unpaired t-test. “*n”* indicates the number of individuals analyzed.

This indicates that, first, heterochromatin maintenance (regulated by DDM1) and maintenance of non-CG methylation (*ddc*) are not fundamental to elicit a defense response against aphid feeding. Second, our result indicates that the roles of KYP in the regulation of H3K9me2 and CHG methylation (Jackson, Lindroth et al., 2002) and/or its uncharacterized role for the maintenance of CHH methylation (Stroud, Greenberg et al., 2013) are an important part of the defense response against aphid infestation. This result correlates with our observed reduction of sRNAs in centromeric and pericentromeric regions (rich in H3K9me2) and the observed changes in CHH and CHG methylation (tightly associated with H3K9me2). Interestingly KYP has been previously associated with the regulation of the defense against geminiviruses (Castillo-Gonzalez, Liu et al., 2015, Raja, Sanville et al., 2008, Sun, Tee et al., 2015) and the maintenance of β-aminobutyric acid (BABA)-induced priming of the salicylic acid (SA)-dependent defense response (Luna, Lopez et al., 2014). In summary, our proof-of-concept analysis indicates that, indeed, mutants in different layers of epigenetic regulation show enhance resistance against aphid settlement.

## DISCUSSION

Organisms monitor environmental conditions and adapt their development according to them. Plants have developed elegant mechanisms of gene regulation adapted to their sessile nature. One of such mechanisms is epigenetic regulation, which could maintain modified transcriptional states through cell division and be reversible once the trigger condition disappears. Although it has been widely proposed that epigenetic regulation is an important part of the stress response, we lack a comprehensive knowledge of the genomic loci that are susceptible to those epigenetic changes and their variability between stresses. Here, we demonstrated that aphid feeding induces changes in the epigenetic regulation of the plant genome and that these changes affect the transcriptional response. Our data suggest that these epigenetic changes are taking place mainly in TEs. We hypothesize that these changes could be important for recruiting TFs that in turn affect the expression of a specific set of defense genes. This will explain while despite having a relatively high number of DMRs (Figure 4), only a very small subset enriched in specific TF-binding motifs are associated with transcriptional changes (Figure 5 and Supplementary Figure 5). An alternative hypothesis to this is that DNA methylation changes are downstream of TF binding, a situation that has been described in human dendritic cells (Pacis, Mailhot-Leonard et al., 2019). Nevertheless, the presence in our analysis of a high number of DMRs without effects at the transcriptional level points against this hypothesis.

Despite their subtle wounding strategy, aphid feeding activates hormonal signals that trigger the reprogramming of the plant transcriptome (Couldridge, Newbury et al., 2007, De Vos et al., 2005, Gao et al., 2010, Kusnierczyk, Winge et al., 2007, Moran, Cheng et al., 2002). Interestingly, the transcriptional changes identified by RNA sequencing show enrichment in genes associated with TF-related activities (Figure 2). These TFs include AR2/ERF and WRKY TFs, which have been associated previously with the transcriptional response against aphid infestation (Foyer, Verrall et al., 2015, Kloth, Wiegers et al., 2016). Counterintuitively, our analysis of the transcriptional and posttranscriptional regulation of TEs during aphid infestation indicated that it is more complex than initially expected from the analysis of the ATH1 data. Our analysis revealed that TEs experience a decrease in the activity of the RdDM pathway translated in a loss of 24 nt sRNAs that leads to their transcriptional reactivation, which is only detectable in deep sequencing experiments via PARE sequencing. This indicates that posttranscriptional regulation of RNA might be an important part of the stress response to aphid feeding. Indeed, posttranscriptional regulation of RNA metabolism is a known regulator of the stress response in eukaryotes (Blevins, Tavella et al., 2019, Harvey, Dezi et al., 2017, Jung, Park et al., 2013, Marondedze, Thomas et al., 2019). These mechanisms could buffer excessive TE transcription to avoid their activity and maintain genome stability during stress-induced transcriptional reprogramming (in our case, loss of 24 nt sRNAs).

The changes of TE activity at the posttranscriptional level prompted us to profile the genome-wide methylation changes under aphid infestation (Figure 4). Our genome-wide analysis of DNA methylation changes induced by aphid feeding show that methylation changes happen primarily at genes (in the CG context) and TEs (in the CHG and CHH contexts). CHH hypomethylated DMRs take place only at epigenetically labile regions characterized by low levels of the repressive histone marks H3K27me3 and H3K9me2 and high levels of the transcriptionally permissive mark H3K18ac. As expected, CHH hypomethylated DMRs are predominantly found at *Helitron* TEs, which are known to influence gene expression (Figure 4). An analysis of the presence of genes in a 4kb window showed us the potential transcriptional changes associated with these DMRs. Between differentially expressed genes associated with DRMs, we found several genes related to the defense response at different levels as *AP2C1* (Schweighofer et al., 2007), *ACS6* (Joo et al., 2008), *SYP122* (Waghmare et al., 2018), *GER5* (Baron et al., 2014), the ethylene response factor *ERF022* and *CCOAMT* (Wang et al., 2017) (Figure 5). Interestingly, 31.25% of the differentially expressed genes associated with CHH DMRs are also differentially expressed in *nrpe1* and/or *ago4* mutants, indicating an influence of the RdDM pathway in the regulation of this response (exemplified by *GER5* in the data showed in Figure 5G). Together with this observation, we found that DMRs associated with differentially expressed genes show an enrichment in binding motifs for certain families of TFs including the AP2-ERF/B3, which has 7 members significantly upregulated upon aphid infestation (Supplementary Figure 5D). These TFs show a modest upregulation in the *nrpe1* mutant and none in an *ago4* mutant, which could be one of the reasons why the transcriptional response differs between aphid infested samples and RdDM mutants. While aphid feeding induces the expression of several TFs, RdDM mutants lack the presence of aphid-induced TFs that would stimulate the defense transcriptional response. As a proof-of-concept, we tested if *Arabidopsis* mutants defective in DNA and histone methylation have a differential susceptibility to aphid infestation (Figure 6). Our analysis indicated that mutations in *Pol IV* and *KYP* show increased resistance to aphid settling, confirming the importance of epigenetic regulation in the response against aphids. In *Arabidopsis* defense genes are located in pericentromeric regions which are densely populated by TEs (Meyers et al., 2003). KYP and PolIV have a known role in the repression of TEs, so we speculate that their lack of function can also facilitate the transcription of genes located in the proximities of TEs. In KYP and NRPD1 mutants, the enhanced activation of defense genes (via transcription or binding of TFs) will explain the increased defense against aphid feeding. Indeed, most of the differentially expressed genes with a proximal CHH DMR identified in our analysis have a TE in the proximities of their regulatory regions (Figure 5).

It is tempting to speculate that together with the downregulation of the epigenetic silencing at DMRs, the observed overexpression of mobile mRNAs and decrease of 24nt sRNAs would trigger transcriptional or posttranscriptional changes on gene expression at distal tissues, other than leaves, including the precursors of the reproductive structures. *Myzus persicae* is known to trigger a transgenerational defense phenotype (De Vos & Jander, 2009). TE silencing is reinforced in the shoot apical meristem (SAM) by the RdDM pathway, what leads to the correct transmission of the right epigenetic states for TEs during vegetative growth (Baubec, Finke et al., 2014). A potential lack of mobile 24nt (Molnar, Melnyk et al., 2010) or 21nt (Dunoyer, Schott et al., 2010) TE-derived siRNAs in the SAM or the reproductive structures, could lead to epigenetic states that could be inherited. Further analysis of the effect of localized stresses on distal tissues and their offspring could share light into the existence of such an elegant overlapping of pathways potentially regulating transgenerational inheritance.

In summary, the evidence presented in our work indicates that changes in epigenetic control are part of the defense response against aphid infestation in *Arabidopsis thaliana*. Intriguingly this response is more complex than previously thought and may involve the interplay between epigenetic and transcriptional regulation. Our work exemplifies the importance of epigenetic regulation in the stress response and the epigenetic plasticity of plant genomes subjected to stress.

## MATERIALS AND METHODS

### Plant and insect material

*Arabidopsis thaliana* (Columbia wild type Col-0, *ddm1-2, ddc, nrpd1a-4* and *kyp-6*) were sown into potting soil (P-Jord, Hasselfors Garden, Örebro, Sweden). At four leaf stage seedlings were selected by uniformity and carefully re-planted into plastic pots (9 × 9 × 7 cm) with one plant per pot at temperature 20–22°C and 70% relative humidity. Plants were grown under L16:D8 light cycle. The light was provided by OSRAM FQ, 80 Watt, Hoconstant lumix, Germany with a light intensity of 220 μmol photons m^-2^ s^-1^. Green peach aphid *Myzus persicae* (Sulzer) was reared in cultures on potted rapeseed plants *Brassica napus* L. under the same climate conditions as the test-plants but in different climate chambers.

### Aphid settling test

An aphid no-choice settling test (Ninkovic, Olsson et al., 2002) was used to investigate aphid behavioral response to different Arabidopsis mutants. One randomly chosen leaf was placed inside a transparent polystyrene tube (diameter 1.5 cm, length 5 cm). The lower end of the tube was plugged with a plastic sponge through which the leaf entered via a slit. Ten wingless *M.persicae* of second to fourth larval instars were placed inside the tube. The upper end of the tube was sealed with nylon net. Leaf of each treatment plant placed inside the tube represented a replicate. The number of aphids settled on the leaf was recorded after 2 h, which is sufficient time for aphids to settle and reach the phloem (Prado & Tjallingii, 1997).

### Tissue for sRNA, RNA, PARE and bisulfite sequencing

5-week-old plants were infested with 40 wingless *M.persicae* of second to fourth larval instars and covered with net cage. After 72 hours all aphids were carefully removed by brush and all *Arabidopsis* rosette leaves were sampled into Falcon tubes extraction bag before being placed in liquid nitrogen. Tissue was collected from all tested mutants. Each frozen plant sample was stored at −70 °C before RNA and DNA extraction.

### DNA and RNA extraction

Total RNA was extracted using TRIzol reagent (Life Technologies) following the manufacturer instructions. mRNA for RNA and PARE sequencing was obtained by purification with the NEB mRNA isolation kit (New England Biolabs). RNA for sRNA library preparation was enriched with the mirVana miRNA Isolation Kit (Life Technologies). Genomic DNA was extracted using the DNeasy Plant Mini Kit (Qiagen).

### Small RNA, RNA and PARE sequencing and analysis

sRNA libraries were produced using the TruSeq Small RNA Sample Preparation Kit (Illumina). Each library was barcoded and sequenced in one lane of an Illumina HiSeq 2000. mRNAs for RNA libraries were isolated using the NEB Magnetic mRNA Isolation Kit (New England Biolabs). RNA libraries were produced using the NEBNext Ultra II Directional RNA Library Prep Kit for Illumina (New England Biolabs). Each library was barcoded and sequenced in one lane of an Illumina HiSeq 2500. PARE libraries were prepared according to Zhai et al (2014). mRNAs for PARE libraries were isolated using the NEB Magnetic mRNA Isolation Kit (New England Biolabs), custom adapters for selecting 5’-P mRNAs and primers from the TruSeq Small RNA Sample Preparation Kit (Illumina) for multiplexing the libraries as indicated in Zhai et al (2014). The resulting sequences were de-multiplexed, adapter trimmed, and filtered on length and quality. sRNAs were matched to the *Arabidopsis* genome, and sequences that did not perfectly align were discarded. Library size was normalized by calculating reads per million of 18‒28 nt genome-matched sRNAs. sRNA and PARE alignments were performed using bowtie (Langmead, Trapnell et al., 2009) with the following parameters –t –v2 that allow 2 mismatches to the alignments. RNA sequencing paired reads were aligned to the Arabidopsis TAIR10 genome using bowtie2 (Langmead & Salzberg, 2012) with default parameters. HTSeq-counts (Anders, Pyl et al., 2014) was used to count reads per gene and the count tables were used in DESeq2 (Love, Huber et al., 2014) to infer significant expression. In htseq-counts for TE analysis the minimum alignment quality value was set to 0 to allow the count of multimapping reads, while this value was set to 10 for analysis of gene expression. Volcano plots were created using ggplot2 (Wickham, 2009). All these tools were used through the Galaxy platform (Afgan, Baker et al., 2018). Heat map for the analysis of microarray data was produced using Heatmapper (Babicki, Arndt et al., 2016).

### Bisulfite sequencing analysis

Adapters and 10 bases from 5’ ends from reads were trimmed using Trimgalore 0.6.1. Clean reads were mapped to the reference genome TAIR 10 using bismark (Krueger & Andrews, 2011) allowing one mismatch per 25 nt seed. Forward and reverse reads were mapped independently. Alignments at the same position were removed using deduplicate_bismark script, including alignments of reads 1 and 2 together. Conversion rate of cytosines were obtained using bismark_methylation_extractor, the first 7 bases from 5’ end and 13 from 3’ end of each read were ignored. The mean conversion rate for the four samples was 99.72%, and the estimated false positive methylation rates were 0.28%. Tile values for genomic DNA methylation were obtained using the Circos: Interval to Tiles pipeline in the Galaxy platform (Afgan et al., 2018). Circular plots were obtained using J-Circos (An, Lai et al., 2015).

### DMR identification

The DMR analysis was carried on with the R package DMRcaller (Catoni et al., 2018), control samples and infected samples were pooled. In order to compare both pools the genome was divided in equal bins of 50 pb size. The DMR were then computed by performing Fisher’s exact test between the number of methylated reads and the total number of reads in both conditions for each bin. The obtained p-values were then adjusted for multiple testing using Benamini and Hochberg’s (Benjamini & Hochberg, 1995) method to control the false discovery. Bins with less than 3 cytosines in the specified context or less than 0.25 difference in methylation proportion between the two conditions or an average number of reads lower than 8, were discarded. Finally bins that were at less than 300 pb were joined.

### Transcription factor binding site prediction

Transcription factor binding site prediction was performed using the plant transcription factor database (http://planttfdb.cbi.pku.edu.cn/). The prediction tool was used for the sequences of the CHH DMRs indicated.

## ACKNOWLEDGEMENTS

Research in the Martinez group is supported by SLU, the Carl Tryggers Foundation and the Swedish Research Council (VR 2016-05410). Sequencing was performed by the SNP&SEQ Technology Platform in Uppsala. The facility is part of the National Genomics Infrastructure (NGI) Sweden and Science for Life Laboratory. The SNP&SEQ Platform is also supported by the Swedish Research Council and the Knut and Alice Wallenberg Foundation.

## AUTHOR CONTRIBUTIONS

Experiment design: RKS, VN and GM. Material contribution: CMP. Performed experiments: MLA, DM, VS and GM. Bioinformatic processing of the data: JLR-V and GM. Data Analysis: GM. Wrote the manuscript: GM. All authors interpreted the data and thoroughly checked the manuscript.

## FIGURE LEGENDS

**Supplementary Figure 1.**
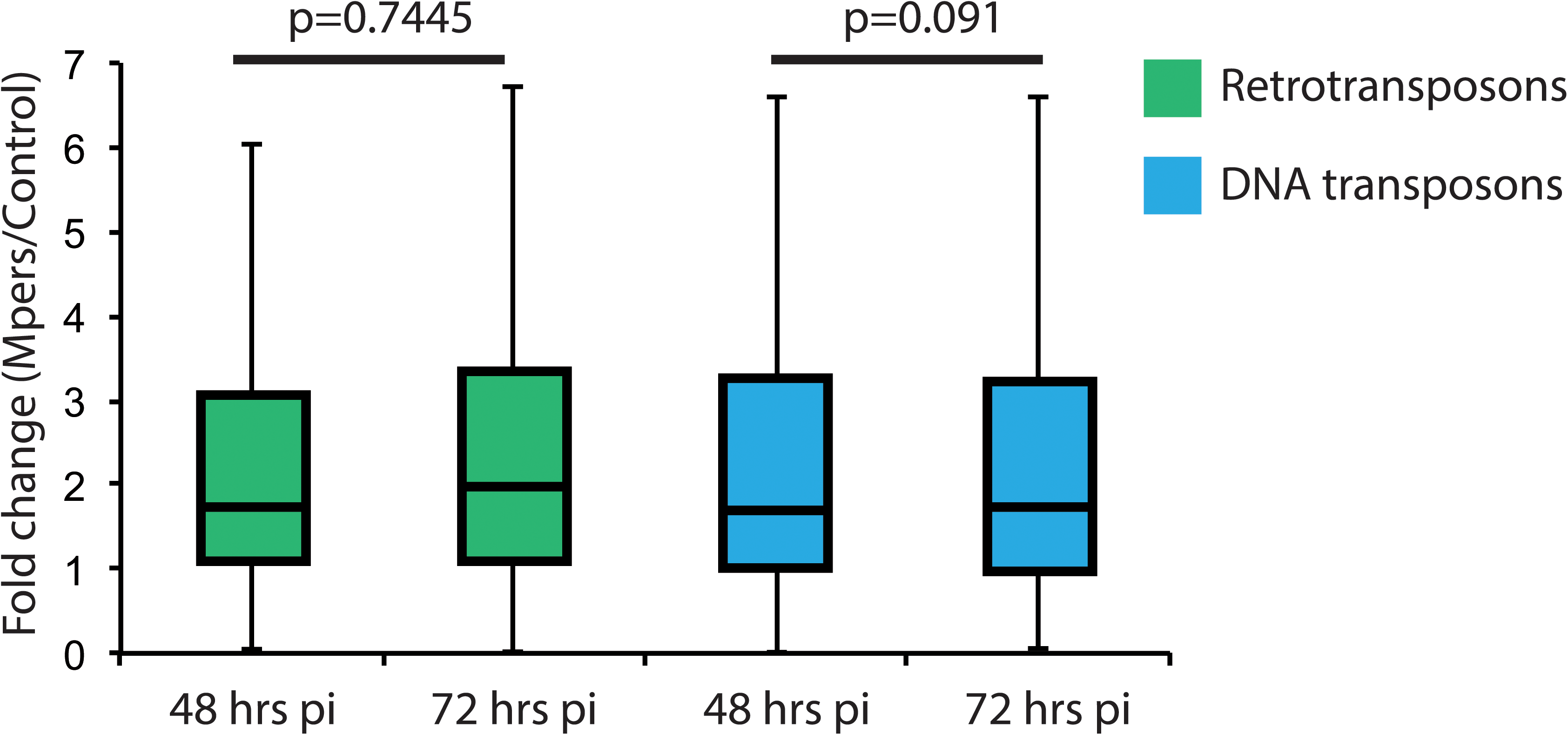
Expression fold change of TEs belonging to different categories in the *Myzus persicae* ATH1 datasets at 48 and 72 hrs pi.

**Supplementary Figure 2.**
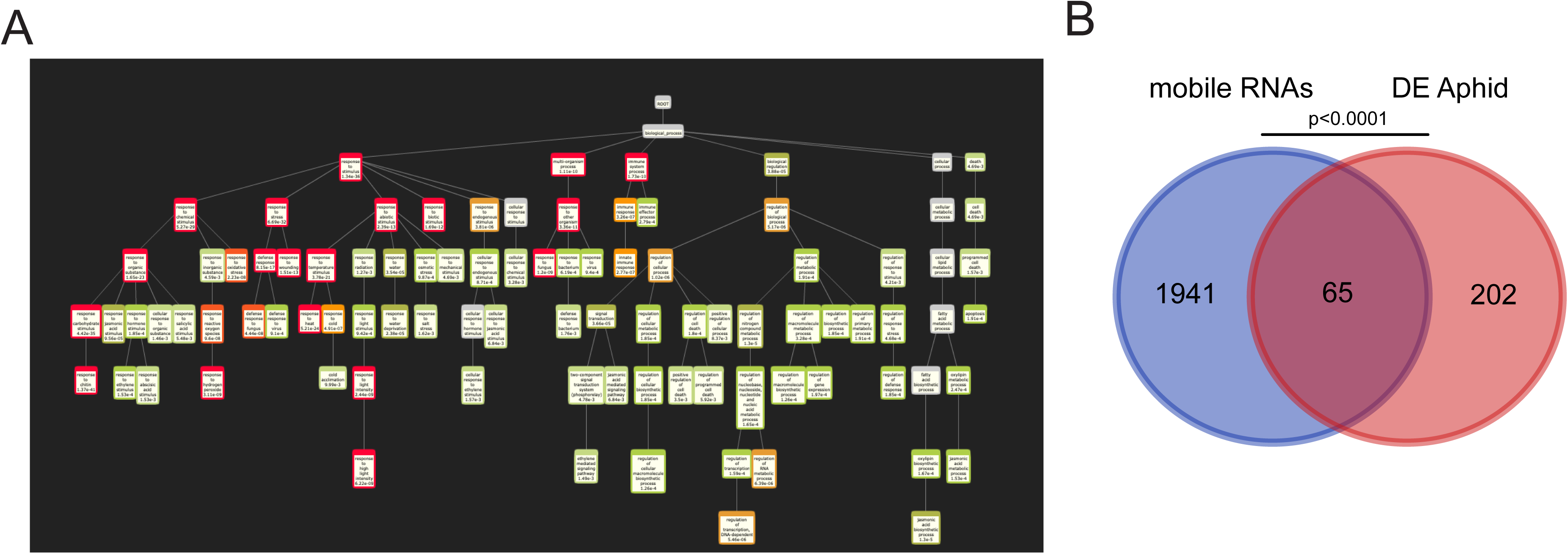
A. Complete biomap of upregulated genes. B. Venn diagram showing the overlap between the whole mobile mRNAs identified in Arabidopsis and the differentially expressed genes during aphid infestation.

**Supplementary Figure 3.**
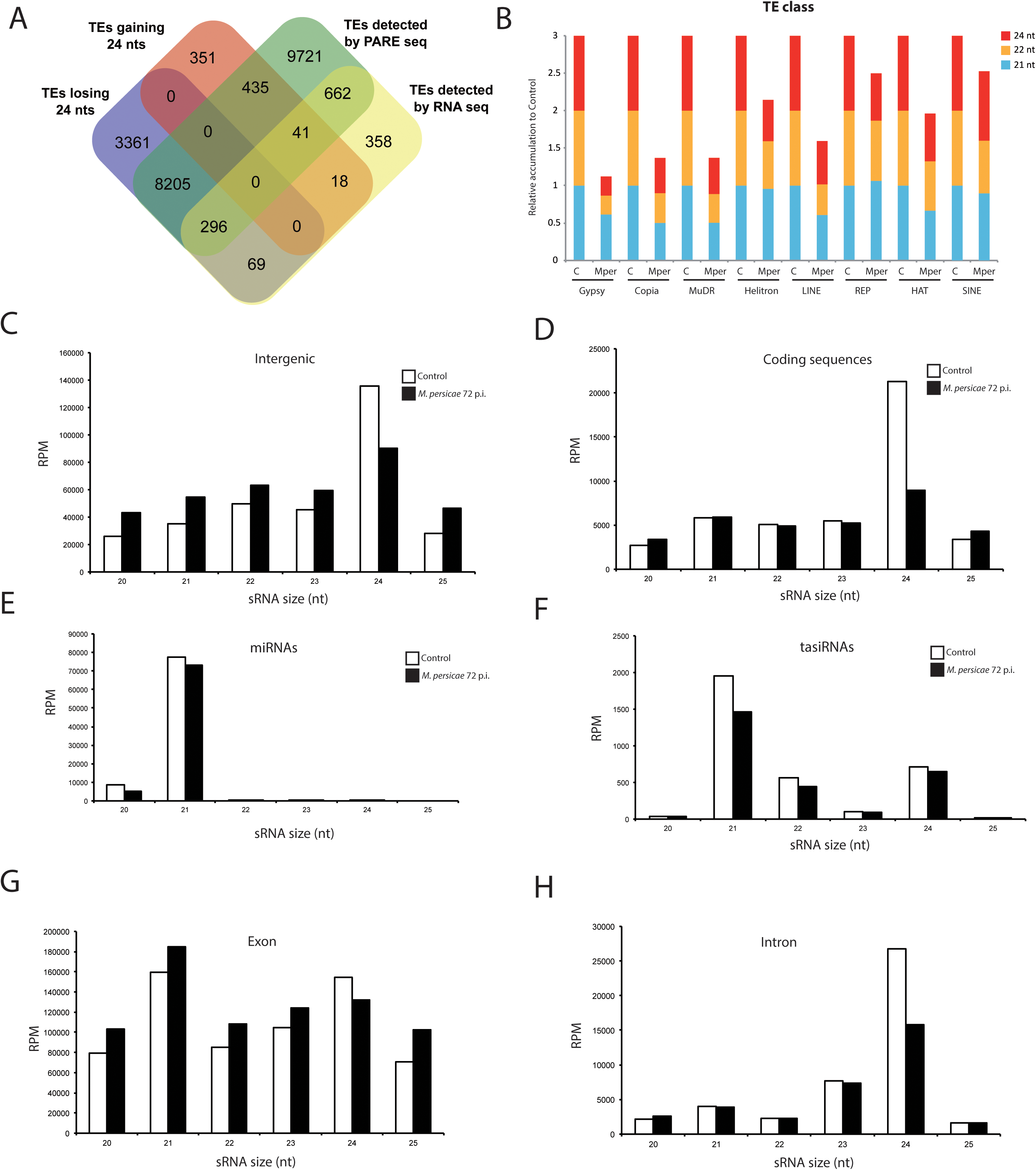
A. Venn diagram showing the overlap of TEs detected by PARE and RNA seq and the TEs that show loss or gain of 24 nts in sRNA sequencing experiments. B. Relative accumulation of 21, 22 and 24 nt sRNAs in aphid infested samples (Mper) relative to control (C) for various TE families. Accumulation values in the control sample were set to one. (C-H) Global sRNAs profiles of control and stressed samples mapping to intergenic regions (C), coding sequences (D), miRNAs (E), tasiRNAs (F), exons (G) and introns (H).

**Supplementary Figure 4.**
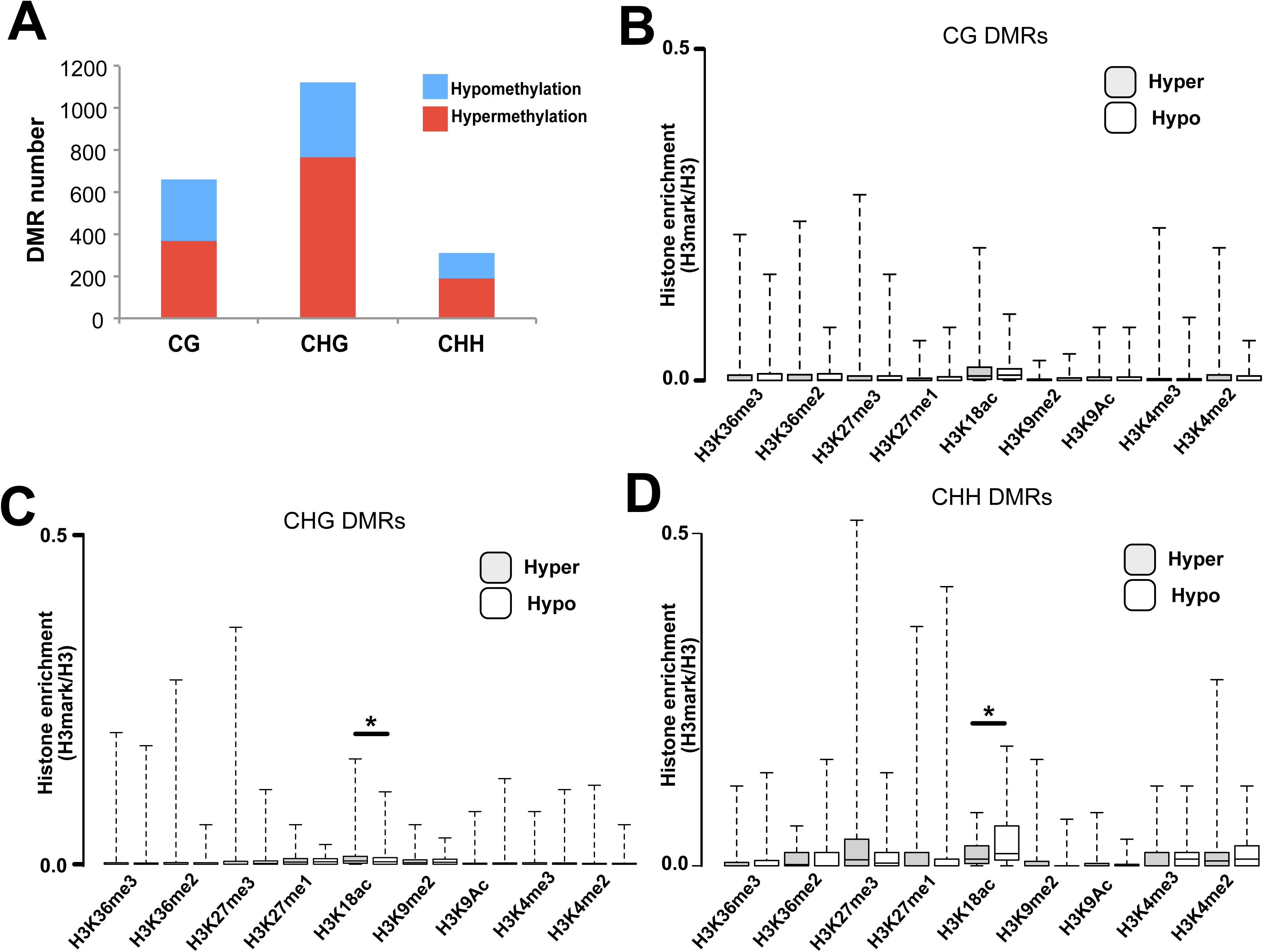
A. Number of hypomethylated (blue) and hypermethylated (red) DMRs present in aphid infested samples for the different DNA methylation contexts. B-D. Histone mark enrichment relative to H3 for hypermethylated and hypomethylated DMRs in the CG (B), CHG (C) and CHH (D) contexts. P-values were calculated using an unpaired t-test.

**Supplementary Figure 5.**
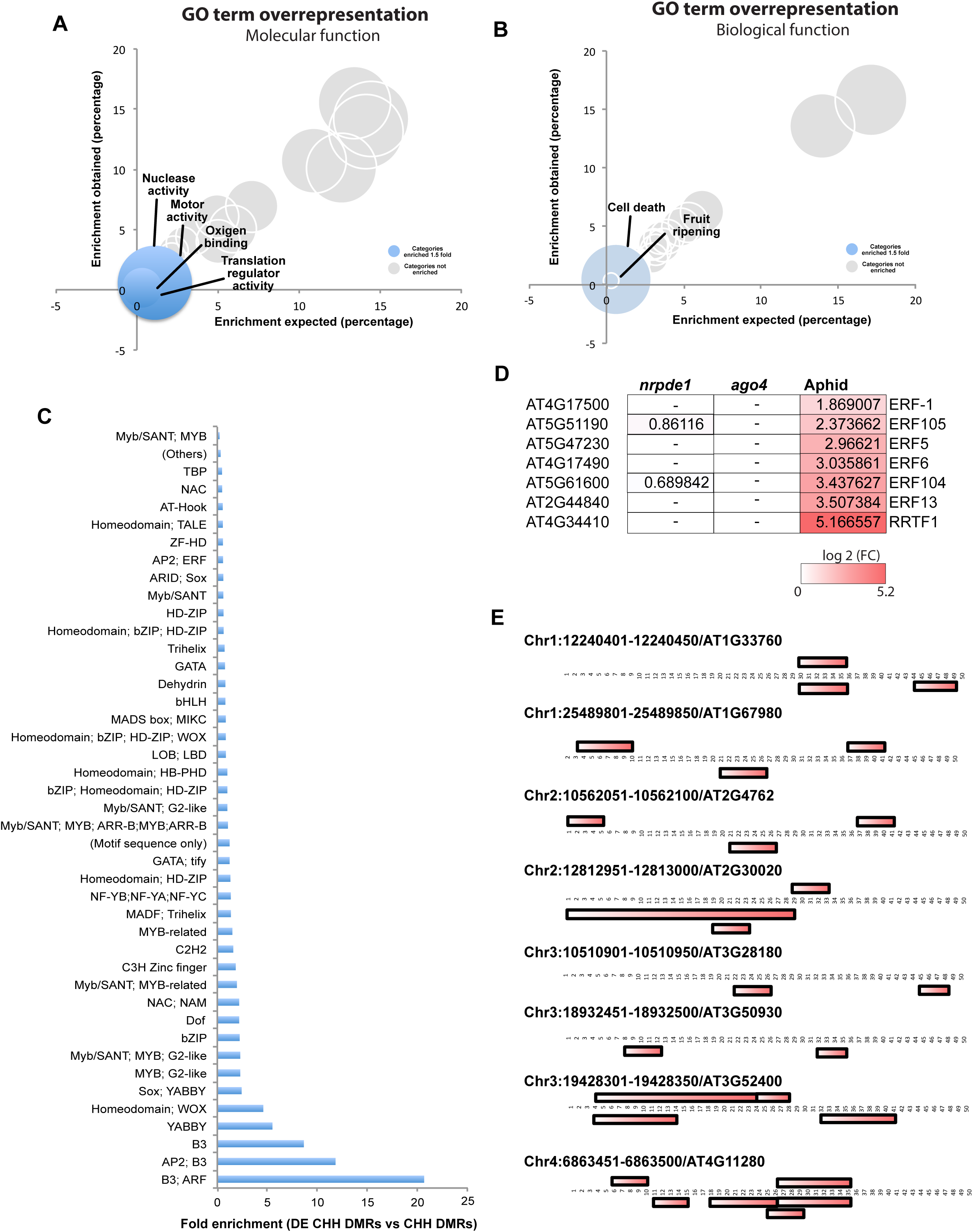
A-B. Bubble graph depicting the GO term overrepresentation test for all genes associated with DMRs grouped by molecular (A) or biological (B) function. Bubbles in blue show GO categories enriched 1.5 fold or more. C. Fold enrichment of transcription factor binding sites at CHH DMRs harboring a differentially expressed gene vs all CHH DMRs. D. Examples of different ERF and ERF/AP2 transcription factors showing upregulation during aphid infestation in *nrpde1*, *ago4* and aphid infested RNA sequencing libraries. Only values of significant differentially expressed genes is shown. E. B3, AP2 and ERF binding sites located at CHH DMRs associated with differentially expressed genes.

